# Structure Insight into Photosystem I Octamer from Cyanobacteria

**DOI:** 10.1101/2021.10.27.465648

**Authors:** Ming Chen, Yujie He, Dongyang Liu, Lijin Tian, Pengqi Xu, Xuan Liu, Yihang Pan, Jun He, Ying Zhang

**Author notes:** To whom correspondence should be addressed: **Jun He** and **Ying Zhang**. These authors contributed equally (Ming Chen, Yujie He).

## Abstract

Diversity of photosystem oligomers is essential to understand how photosynthetic organisms adopted to light conditions. Given by the structural and physiological significance, the assemblies of PSI supercomplex is of great interest in both chloroplast and cyanobacteria recently. In this study, two novel photosystem I supercomplexes were isolated for the first time from the low light incubated culture of filamentous cyanobacterium *Anabaena* sp. PCC 7120. These complexes were defined as PSI hexamers and octamers through biochemical and biophysical characterization. Their 77K emission spectra indicated that the red forms of chlorophylls seemed not to be affected during oligomerization. By cryo-EM single particle analysis, a near-atomic (7.0 Å) resolution structure of PSI octamer was resolved, and the molecular assemblies of stable PSI octamer was revealed.

## INTRODUCTION

Photosystem I (PSI) is a light-driven plastocyanin: ferredoxin oxidoreductase with a quantum efficiency close to 100%, probably the most efficient photoelectron converter in nature (Amunts and Nelson, 2009). The function of this bio-macromolecule is highly conserved among cyanobacteria, algae, and plants, yet carries complicated structural diversities, especially in cyanobacteria. It has been acknowledged for years that chloroplast PSI-LHCI complex can only form monomeric state. And this classic theory was challenged by an observation of spinach thylakoid membrane with atomic force microscopy (AFM), in which PSI-LHCI complexes tense to be assembled each other to form a dimeric supercomplex in dark condition (Wood et al., 2018). And an atomic-resolution (2.97 Å) structure of novel dimeric PSI-LHCI supercomplex was resolved in low light adapted green alga *Chlamydonomas reinhardtii* very recently (Naschberger et al., 2021). Things are more complicated in cyanobacteria, in which the PSI complex is assembled as a trimer in most species (Boekema et al., 1987, Kruip et al., 1994, Jordan et al., 2001), meanwhile, the dimer and tetramer have been identified in some heterocyst-forming cyanobacteria and their close unicellular relatives (Watanabe et al., 2014; Li et al., 2014; Li et al., 2019, Zhang et al., 2019, Chen et al., 2020). Recently, high-resolution structures of PSI monomers with full physiological function were also resolved in some species of unicellular cyanobacteria (Netzer-El et al., 2018, Çoruh et al., 2021). All these studies have indicated that the dynamic oligomeric states of PSI complex are essential to help photosynthetic organism adapting the changing environments, especially under stress conditions, such as limited light energy supplying.

Moreover, by using blue-native PAGE, Li and co-workers observed a possible PSI hexamer in *Chroococcidiopsis* sp. PCC 6712 upon thylakoid membrane extraction (Li et al., 2019). And before that, a projection of two associated PSI tetramers was spotted with single particle analysis in filamentous cyanobacterium *Anabaena* sp. PCC 7120 (here after *Anabaena* 7120) (Watanabe et al., 2014). Due to the key role of PSI oligomer states in understanding the photosynthesis mechanism, molecular evidences and structure insights to those higher-ordered PSI oligomer complexes are of significant interests and highly desired now. Here, two novel oligomer states of PSI complex were isolated from the low light incubated culture of *Anabaena* 7120. The pure molecules were set into biophysical analysis, and the existence of higher-ordered PSI supercomplexes in cyanobacteria was ultimately demonstrated. By employing cryo-EM single particle analysis, one of these PSI supercomplexes was subjected to high resolutions structure determination, in which a novel molecular interface for PSI complex assembling were detected.

## RESULTS AND DISCUSSION

### Discovery of higher-ordered PSI complexes

Blue-native PAGE has been recognized as a powerful tool to investigate thylakoid membrane complexes (Wittig et al., 2006, Sari et al., 2011). The bands corresponding to complexes of high molecular weight suggest the existence of higher-order oligomeric states of photosynthetic membrane complexes currently undefined, especially for PSI assembled hetero- or homo- complexes (Sari et al., 2011; Kouril et al., 2018; Zhang et al., 2010).

As shown in figure 1a_1_, two previous undescribed complexes (supercomplex 1 and 2), both larger than the PSI-tetramer (1400 kDa), were extracted from *Anabaena* 7120 using 1% (w/v) β-DDM and subsequently separated on BN-PAGE gel. The naturally green colour of the bands indicates that these are chlorophyll-containing complexes, most likely the PSI-assembled complexes. To verify this hypothesis, the complexes were separated and harvested through sucrose gradient ultracentrifugation. The fraction scattering in sucrose gradients (Figure 1a_2_) is highly consistent with the gradient gel results in the BN-PAGE experiment (Figure 1a_1_). The fractions corresponding to supercomplex 1 and supercomplex 2 were collected and subjected to low temperature (77 K) fluorescence spectra measurement, and the PSI tetramer fraction was also harvested for comparison. As shown in Figure 1b, the emission spectra of supercomplexes 1 and 2 peaked at ~730 nm, largely overlapping with the one of PSI tetramer. Their emission spectra showed that the lowest energy level of quantum yield of chlorophylls in PSI remained ~730 nm, indicating that the red forms of chlorophylls were not energetically altered. This fluorescence spectral characteristic strongly suggests that these two complexes are both PSI homogenous supercomplexes (Watanabe et al., 2014, Lamb et al., 2018, Chen et al., 2020).

**Figure 1.**
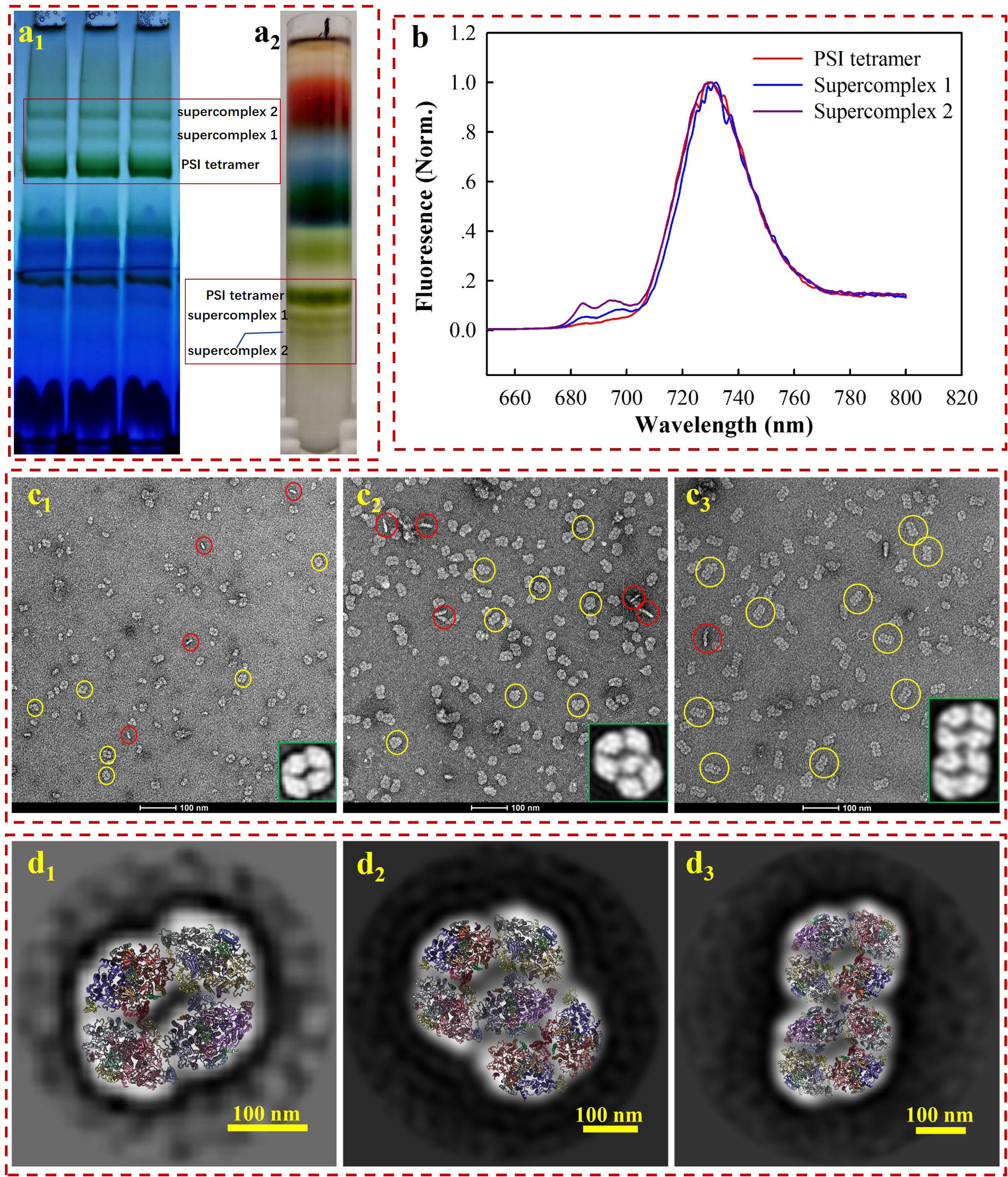
Biochemical (a) and biophysical (b, c, and d) characterisation of the PSI tetramer, supercomplex 1, and supercomplex 2. a_1_-membrane crude extracts were loaded on BN-PAGE at a total amount of 9 to 12 mg chlorophyll a for each lane; a_2_-membrane crude extracts were loaded on sucrose gradient ultracentrifugation for the separation of PSI tetramer and supercomplexs. b-low temperature (77k) fluorescence emission detection of PSI tetramer (red line), supercomplex 1 (blue line), and supercomplex 2 (purple line). c-Negative staining electron microscopy image of PSI tetramer (c_1_) supercomplex 1 (c_2_) and supercomplex 2 (c_3_). The yellow and red circles indicate the top and side views, respectively. The enlarged image in the right corner shows the 2D averages of the most contributed particles. d-The model fitted 2D average maps of PSI tetramer (d_1_) supercomplex 1 (d_2_) and supercomplex 2 (d_3_).

For a closer look at the molecules, the complexes were subjected to negative staining electron microscopy (EM) single particle analysis. The PSI tetramer, supercomplex 1, and supercomplex 2 were stained with 0.75 % (w/v) uranyl formate and observed by transmission electron microscopy (TEM). While the homogeneity of particles from all three samples is acknowledged (figure 1c_1_, 1c_2_ and 1c_3_), the remarkable diversity in particle size from different samples is also revealed by negative staining EM. We subsequently using single particle analysis (SPA) approach to check the oligomeric states of various samples. Using the high-resolution structure of PSI tetramer as reference (Zheng et al., 2019; Kato et al., 2019; Chen et al., 2020), particles in micrograph figure 1c_1_ are tilted views of the PSI tetramer. Over 90% of the particles presented a top-view (yellow circles) orientation parallel to the thylakoid membrane plane, while about 9.5% of them are side views (red circles). The contribution of the top and side views is consistent with previously report (Watanabe et al., 2014). For the samples of supercomplex 1 and supercomplex 2, the particle morphology in the images indicated that these proteins are PSI higher-ordered oligomeric complexes (figure 1c_2_ and 1c_3_).

To further characterize the molecular architectures, three datasets of 16, 37, and 18 micrographs were collected for the PSI tetramer, supercomplex 1, and supercomplex 2, respectively, from which 359, 2518, and 822 particles were manually picked. Particles were then used for pre-processing and 2D classification, and the most dominant averages are shown as the enlarged part in the bottom right corner of figure 1c_1_, figure 1c_2_, and figure 1c_3_. From these closer views, it is strongly suggested that supercomplex 1 is a PSI hexamer consisting of a tetramer and a dimer, while supercomplex 2 is a PSI octamer formed by two assembled tetramers, which agree with the previous results (Watanabe et al., 2014). This conclusion is supported by a rigid fitting of PSI tetramer and dimer models over the 2D map of the complexes, respectively (figure 1d). By comparing to the PSI tetramer, the PSI hexamer can be identified as a complex arranged with a tetramer and dimer with substantial confidence. However, it is ambiguous to identify the interface for the linkage of the PSI tetramer and dimer in such low-resolution map. Furthermore, from the fitted model (figure 1d_2_), the PsaL-PsaA, PsaK-PsaK, and PsaM-PsaF interfaces should be addressed in further studies providing high resolution details. PSI octamer revealed in figure 1d_3_ is a supercomplex assembled though ‘side by side’ arrangement of two tetramers, the fitted model reveals the possible interfaces of PsaA-PsaF, PsaK-PsaK, and PsaB-PsaK. Studies providing high-resolution structure would also be appreciated to investigate the detailed molecule basis for the intra- and inter- connection of these supercomplexes.

### Cryo-EM structure of PSI octamer complex

In our experiments, the in vitro safety and stabilities of PSI octamer is better than the hexamer, for this reason, a near-atomic resolution structure of stable PSI octamer was resolved by cryo-EM single particle analysis in the present study. For maximal harvests and minimal destroy to PSI octamer, Native-PAGE based purifying method was applied for sample purification. The whole processes for concentrated pure samples preparation were completed in only six to eight hours, and the fresh molecules were than frozen immediately for cryo-EM single particle analysis. Protein samples with a concentration of approximately 3 mg/mL were loaded onto the grids for frozen and quality evaluation. Initially, very few particles could be observed in the hole of the ice layer, while most of them clumped together around the carbon edge (Figure 2a_1_). To help particles move into the hole, an extra detergent, fos-choline 8, was added to the sample immediately just before the blotting. A significantly improved distribution of particles in the holes was obtained, with approximately 30 particles per micrograph (Figure 2a_2_). However, the presence of random orientations is a challenge when considering the surface charge of the complex. By optimising the wait time between sample loading and blotting, we improved the distribution of the side view by approximately 10% (figure 2a_3_). The preferential orientation issue is biological specimen dependent, and many options may proceed case by case (Han et al., 2020, Sorzano et al., 2021, Tan et al., 2017). If the particles are two linked PSI tetramer, then the preferential orientation is relatively inevitable as the hydrophilic area of top view is as large as around 1000 nm^2^, while it is less than 200 nm^2^ in side view (supplementary figure 1). And top views have much higher probability to be absorbed and face to the air water interface when sample being blotted and frozen (Noble et al., 2018).

**Figure 2.**
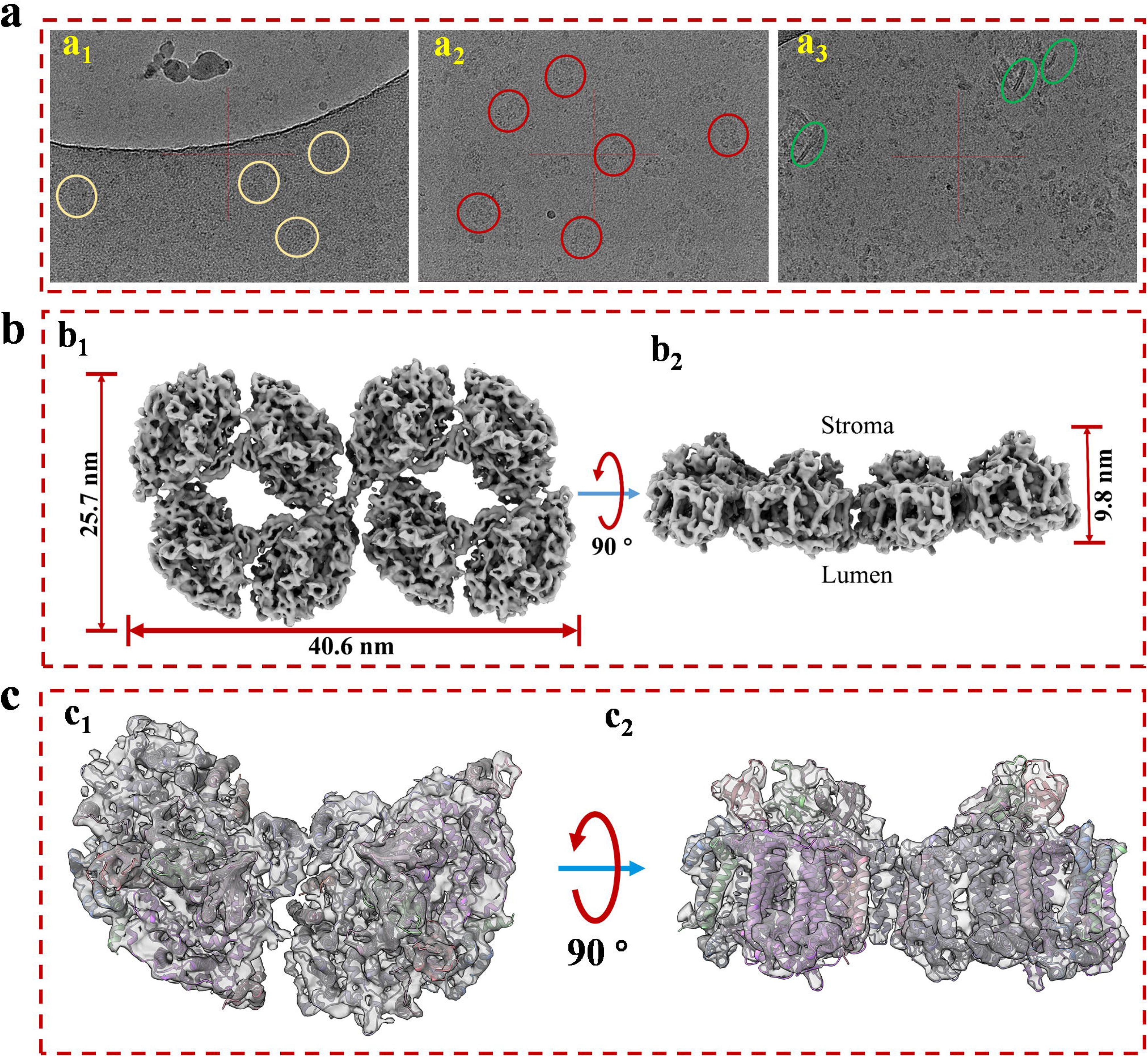
The cryo-EM sample prep optimization and structure determination of PSI supercomplex 2. a-Optimization of cryo-EM sample. a_1_-The protein particles were blotted immediately after sample loading; a_2_-Detergent fos-coline-8 were supplemented into protein sample at final concentration of 1.0 mM around 5 seconds before blotting; a_3_-Additional 3 seconds waiting time were introduced between the loading and blotting of fos-choline-8 mixed sample. b-Top (b_1_) and side (b_2_) views of the refined 3D density map of PSI supercomplex 2. c-Model and map fitting for structure comparison of cyanobacterial PSI dimer and the supercomplex. Top (c_1_) and side (c_2_) views of model fitting of PSI dimer into the density map. The model used for fitting is a published PSI dimer structure (PDB code: 6K61) from *Anabaena* sp. PCC 7120.

In data collection stage, 23491 movies were collected from 17 datasets using a 200 kV microscope equipped with a Gatan K3 camera. As revealed in supplementary figure 2, in total 184882 particles were extracted during data processing, from which 124460 good particles were randomly separated into 10 datasets and screened for 3D reconstruction and structure determination respectively in cryoSPARC. Reconstruction of this complex proceeded without applying symmetry (C1) from homogenous particle subsets. Non-uniform refinement from 12121 particles of subset 5 generated a map at overall resolution of 7.0 Å and highest local resolution of 6.0 Å (supplementary figure 3). As shown in figure 2b, the molecular size of PSI octamer are 40.6 nm, 25.7 nm and 9.8 nm on length, width, and height respectively. The overall shape of the map strongly suggested that this megacomplex is a PSI oligomer synclastic arranged by two linking tetramers side by side, which first proved that the complex is a PSI octamer structurally. To verify this conclusion a PSI dimer model was fitted into a part of the map (figure 2c_1_ and 2c_2_). The secondary structures in both of the transmembrane and out membrane region (lumen and stroma sides) can be fitted perfectly into the densities.

Interestingly, a density of interface for linkage was objectively reconstructed in this map. As shown in supplementary figure 4, the density (red frame) is consistent with the threshold levels, and the firmness is stronger than the interfaces that forms dimers (green frame) and tetramers (light blue frame). It is indicated that the newly identified PSI octamer is a physiologically relevant megacomplex rather than a random aggregation during sample preparation. This solid evidence was then obtained from the biochemical characterizations of PSI octamer from cells growing in changing physiological conditions. As shown in supplementary figure 5, by native-PAGE characterization of cells that incubated in different light conditions, the PSI octamer is a low light responded supercomplex in *Anabaena* sp. PCC 7120. The result agrees with the conclusion from Wood’s work published in 2018, in which tighter packing of PSI complexes were detected in the dark (Wood et al., 2018). To find the possible interfaces for the assembly of two tetramers, separated PSI tetramer model were fitted rigorously into the map with ChimeraX program. As shown in figure 3, an interaction between PsaK subunits from two linked PSI tetramers was found in the fitted model (figure 3a). As revealed in figure 3b, two residues Lys52 and Phe53 of PsaK from tetramer 1 (T1) interacted with the same residues from tetramer 2 (T2) with each other. This novel interface should be the major contribution to stabilize PSI octamer. In summary, the map is qualified to be applied to perform rigid-body fitting with an atomic model when more structural details are required to be revealed. On the other hand, this map is reconstructed with datasets collected from the 200 kV microscope, and higher resolution is expected by using a zero-loss energy filter equipped detector, and the resolution can also be theoretically significantly improved with a 300 kV microscope.

**Figure 3.**
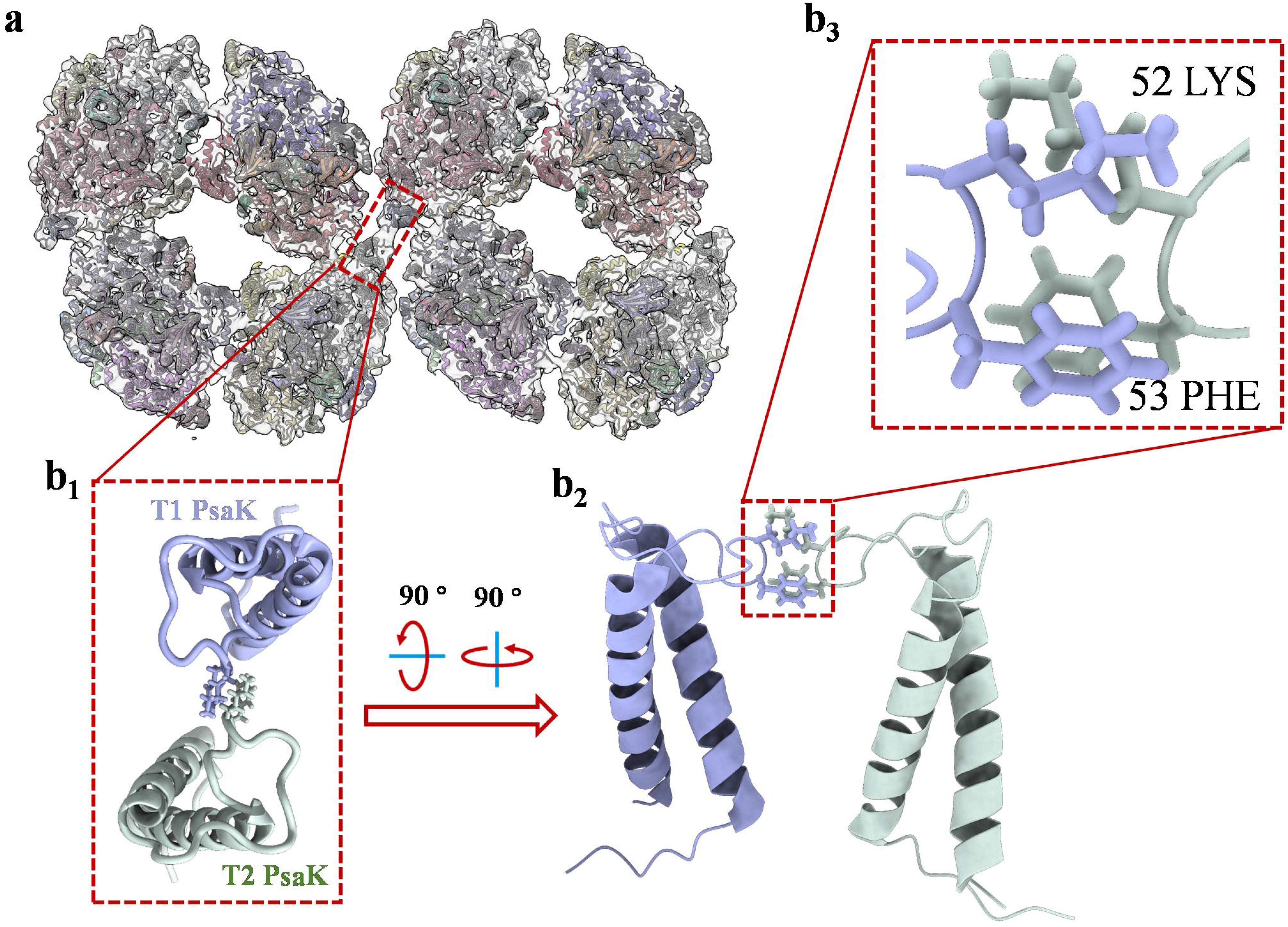
Structure details of PsaK-PsaK interaction in PSI octamer supercomplex. a-Overall view of the interface location in the complex (top view from stromal side). The original structure applied for fitting is PSI tetramer model (PDB code: 6TCL) from *Anabaena* sp. PCC 7120. b-Close-up view of structural details indicated in figure 3a. b_1_-The enlarged top view of two PsaK subunits from the assembled PSI tetramers (T1 and T2), the interacted amino acids were shown as stick form. b_2_-Tilted side view of interacted PsaK subunits. b3-The focus view of intact amino acids Lys52 and Phe53 in PsaK subunits.

Higher-ordered PSI oligomers are physiologically relevant megacomplexes rather than random aggregations during sample preparation. For instance, in this study, the PSI octamer was strongly suggested as a low light responded supercomplex in *Anabaena* sp. PCC 7120 with the results of supplementary figure 5. Similar conclusion was presented by Wood’s observation through atomic force microscopy, in which tighter packing of chloroplast PSI complexes are more abundant in the dark (Wood et al., 2018). And the super high-resolution structure of such tight packed chloroplast PSI oligomer was resolved very recently (Naschberger et al., 2021). Novel type of interfaces for connection between PSI monomers were structurally detected and genetically solidified. PSI assembled heterogenous or homogenous supercomplexes are often detected biochemically in previous studies (Sari et al., 2011; Kouřil et al., 2018; Zhang et al., 2010); here, we isolated two supercomplexes and set up biophysical characterisations of the molecules. Taking the evidence from low-temperature fluorescence and negative staining EM, we concluded that these two complexes are PSI hexamer and PSI octamer, respectively. To our knowledge, this is the first study to isolate such higher-ordered PSI complexes in photosynthetic organisms, paving the way for better understanding the molecular architecture and physiological functions of each type of photosystem oligomers. And cryo-EM structure of PSI octamer revealed a novel molecular interface (PsaK-PsaK) for PSI assembling in photosynthetic organisms.

## MATERIALS AND METHODS

### Specie and growth conditions

Filamentous cyanobacteria *Anabaena* 7120 (FACHB-418), were purchased from Institute of Hydrobiology, Chinese Academy of Sciences. Cyanobacterial cells were cultured at 20°C to 30°C in BG-II (+N) media. The liquid culture in flask were incubated under illumination with fluorescent lamps (5-50 μE m^−2^s^−1^).

### Membrane protein complexes solubilization

For membrane complex extraction, 150 mL liquid cultures at the logarithmic phase (A_730_=1.0 to 1.3) were harvested at 4000 g centrifugation under 4°C, and cells was washed once in 15 mL pre-cooled buffer A (50 mM HEPES, 10 mM MgCl_2_, 5 mM CaCl_2_ and 15 mM NaCl). Cells were then resuspended in 10 mL buffer A and treated with a cycle of 15 minutes frozen in −80 °C refrigerator and 10 minutes melting in room temperature. Treated cells were harvested at 4000 g centrifugation under 4°C and resuspended in 1.5 to 3 mL buffer A, and 10% (w/v) of β-DDM were supplemented to the given final concentration for membrane solubilization and complex extraction. The mixture was incubated on ice for 30 minutes, and insolubilized materials were removed by 14,000 g centrifugation under 4°C for 15 minutes. The supernatant in which membrane complexes resolved were harvest and store at 4°C, or directly proceeded for BN-PAGE analysis.

### Isolation of membrane supercomplexes

To isolate membrane supercomplexes, solubilized supernatant was collected and subjected to sucrose gradient ultracentrifugation (10% to 50% (w/v) sucrose in buffer A with 0.03% (w/v) digitonin) at 35000 rpm (sw41 Ti rotor, Beckman Coulter) for 16 hours at 4℃. The band containing PSI tetramers and lower bands were collected with a syringe. The collected fractions were buffer exchanged using a Milipore concentrator with 100 kDa molecular weight cutoff into buffer A with 0.03% (w/v) digitonin to remove the sucrose and to concentrate the sample.

### Blue-Native PAGE and reextraction

BN-PAGE was performed in a cold room or a large ice bucket to maintain the low temperature (4°C) during electrophoresis. A Bio-Rad Mini-PROTEAN Tetra Cell (1658006) system was used for mini gel casting and electrophoresis running. The reagents for gel casting, the anode and cathode native PAGE buffers were prepared according to Witting’s protocol (Witting et al., 2006). A gradient generator was used to prepare the acrylamide gradient gel. In this study, 3%–10% (w/v) acrylamide gel was selected according to the large molecular weight of megacomplexes. In most cases, 4%–13% (w/v) gels are feasible to separate proteins with molecular weights ranging from 10 kDa to 3,000 kDa, and 3%–13% (w/v) gels are appropriate for separating 10 kDa to 10,000 kDa protein molecules. Electrophoresis power was generated using a Bio-rad PowerPac TM Basic Power Supply (1645050). In the first stage, cathode buffer B was used, and the power supply was set to 135V for electrophoresis. Move into stage two by exchanging cathode buffer B to cathode buffer B/10 when the samples were moved into a separate gel (approximately 30 min). The running was maintained at 135V for another 1.5 to 2 hours until acceptable separation efficiency was obtained.

For the reextraction of proteins, target bands were excised immediately after electrophoresis and stored in an ice-cooled Eppendorf tube. Gel bands were manually homogenised with a glass rod directly into the tube to minimise the loss of protein samples. To prevent heat damage to the protein complex, 10 cycles of 30 s on and 60 s off were programmed for homogenisation. The gel slurry was then mixed with reextraction buffer (buffer A with 0.03% digitonin), and the gel debris was removed by centrifugation at 14,000 × g for 15 min at 4°C. The supernatant was subjected to a second run of BN-PAGE for purity and integrity check, and concentrated to qualified concentration with a 100 kDa cut off Millipore membrane filter (UFC5100BK) for single particle analysis.

### Cryo-EM and Image Processing

Three microliters of purified PSI supercomplex was applied onto glow-discharged holey carbon grids (Cu Quantifoil R1.2/1.3) at a protein concentration of approximately 2.5 mg/ml, prior to vitrification using a Vitrobot MKIV (3.0 s blot 5, 4 °C, 100% humidity). The images were collected in 17 different sessions on 200 kV FEI Titan Krios electron microscope (50 μm C2 aperture). A Gatan K3-Summit detector was used in counting mode at a magnification of 45,000 (yielding a pixel size of 0.88 Å), and a dose rate of 30 electrons per pixel per second. Exposures of 1.618 s (yielding a total dose of 63 eÅ^−2^). SerialEM was used to collect a total of 23491 images, which were fractionated into 27 movie frames with defocus values ranging from 1.5 to 2.5 μm. All datasets were processed separately by using the same procedure. Collected micrographs were corrected for local-frame movement and dose-filtered using Motioncor2 (Zheng et al., 2017). The contrast transfer function parameters were estimated using Gctf (Zhang, 2016). In total, 184882 particles were autopicked using warp and subjected to reference-free two-dimensional class averaging in Relion 3.0.8 (Zivanov et al., 2018). After 2D classification, 12k good particles were generated, in which top-views were divided into 10 subsets. Each subset was combined with the side-view separately for the 3D map refinement in cryoSPARC (Punjani et al., 2017), yielding a map at an overall resolution of 7.0 Å. No symmetry was applied during the final processing, as it did not improve the map. The reported resolutions are based on 3D refinement by applying 0.143 criterion on the FSC between reconstructed half-maps. All figures were generated using PyMOL (DeLano et al., 2002) and Chimaera X (Goddard et al., 2018). A local resolution map was generated using the cryoSPARC.

### Negative staining EM and data processing

The purified PSI tetramer and supercomplexes were adjusted to 0.15, 0.19 and 0.27 mg/ml protein respectively. In all cases, 3 μl of the protein solution was loaded on the glow-discharged copper grid. The excess protein solution was blotted by filter paper. The grid was washed twice with distilled water, and then stained with 0.75 % (w/v) uranyl formate for 30 s. The stain solution was blotted and the grid was dried in air. The grids were examined using a Tecnai G2 Spirit electron microscopy under 120 kV accelerating voltage. Micrographs were recorded with CCD camera at a normal magnification of 49,000× with variation of defocus values from −1.5 to −2.5 μm, resulting in a pixel size of 4.3 Å at specimen level. In total 34, 16 and 18 images were manually collected for PSI dimer, tetramer and supercomplex. 359, 2518 and 822 particles were manually picked for PSI tetramer, and supercomplex in Relion 3.0.8. The selected particles were proceeded to a reference-free two-dimensional class averaging using Relion 3.0.8.

## ACKNOWLEDGEMENT

This work was supported by China Postdoctoral Science Foundation (Grant No. 2020T130747) and by 100 Top Talents Program of Sun Yat-sen University (Grant NO. 392009).

## AUTHOR CONTRIBUTIONS

M.C., X.P. and Y.Z. designed the research; Y.H, M.C. and D.L. performed the research and data curation under supervision of L.T. and J.H.; M.C. wrote the manuscript with adding from Y.H., Y.P. and D.L.; Y.Z., L.T and J.H. analysed the data and edit the manuscript; J.H. and Y.Z. supported the research with funding acquisition. All authors read and approved the manuscript.

## CONFLICT OF INTETRESTS

The authors declare no conflict of interest.

