## Supplemental figures for "Structure Insight into Photosystem I Octamer from Cyanobacteria"

Authors’ affiliations: 1, The Seventh Affiliated Hospital, Sun Yat-sen University, Shenzhen, China. 2, Center for Cell Lineage and Development Guangzhou Institutes of Biomedicine and Health, Chinese Academy of Sciences, Guangzhou, China. 3, Center for Cell Fate and Lineage (CCLA), Bioland Laboratory (Guangzhou Regenerative Medicine and Health Guangdong Laboratory), Guangzhou, China. 4, Photosynthesis Research Centre, Institute of Botany, CAS., Beijing, China.

#These authors contributed equally.

**Supplementary Information**

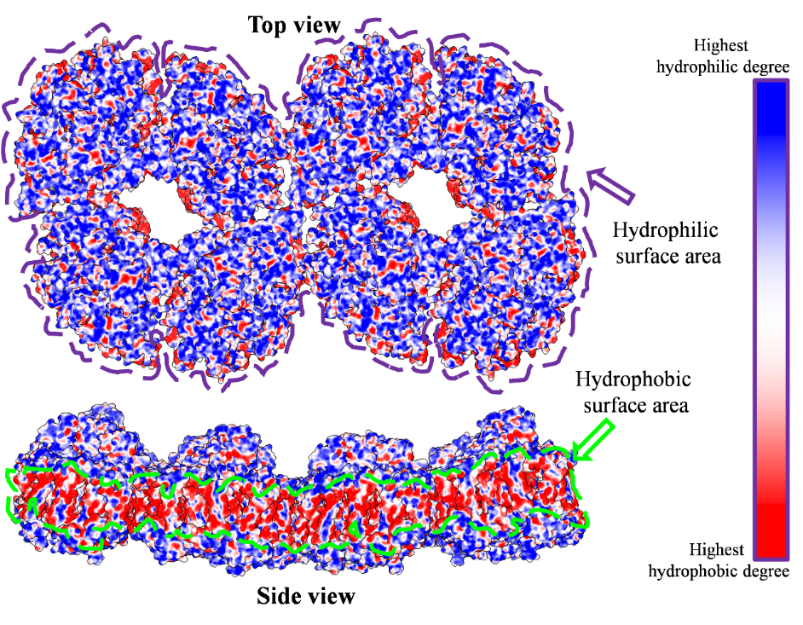

**Supplementary figure 1.** Hydrophobic area analysis of PSI octamer model. The hydrophilic area shown in the figure (Top view) is the luminal side surface of PSI complex. The hydrophobic surface area shown in the figure (Side view) is the transmembrane region. Blue color represents the most hydrophilic area and red color represent the most hydrophobic area of the complex.

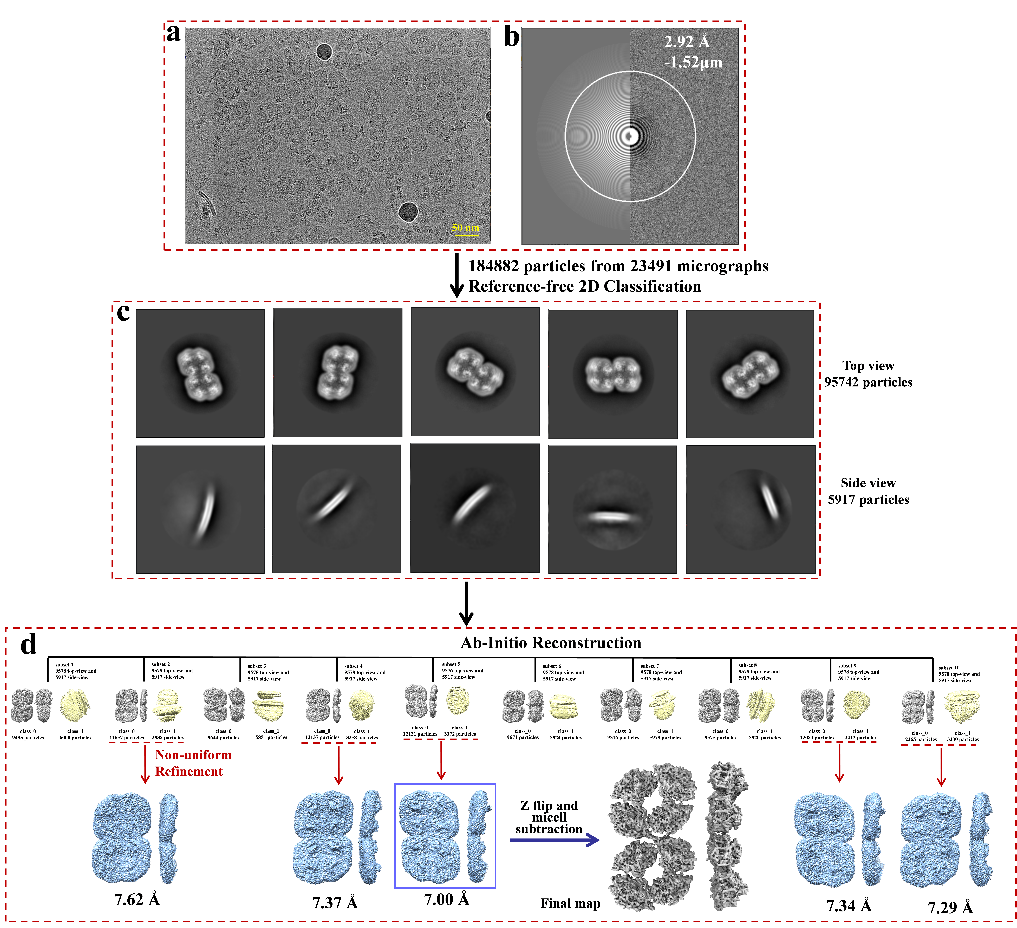

**Supplementary figure 2.** Cryo-EM image data processing. a-Representative micrograph. b-CTF indicating max resolution. c-Representative 2D classes. d-3D reconstruction and refinement.

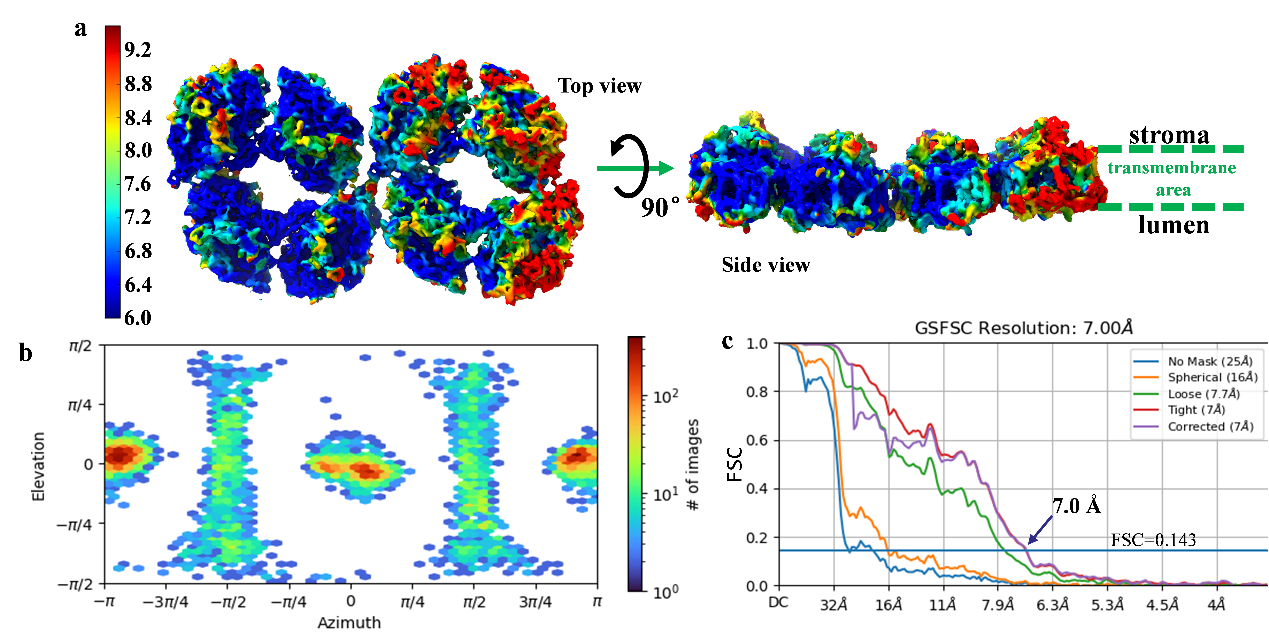

**Supplementary figure 3.** Resolution estimation of reconstructed map. a-Local resolution map of PSI supercomplex; b-Direction distribution of particles for 3D reconstruction; c-GSFSC resolution estimation curve.

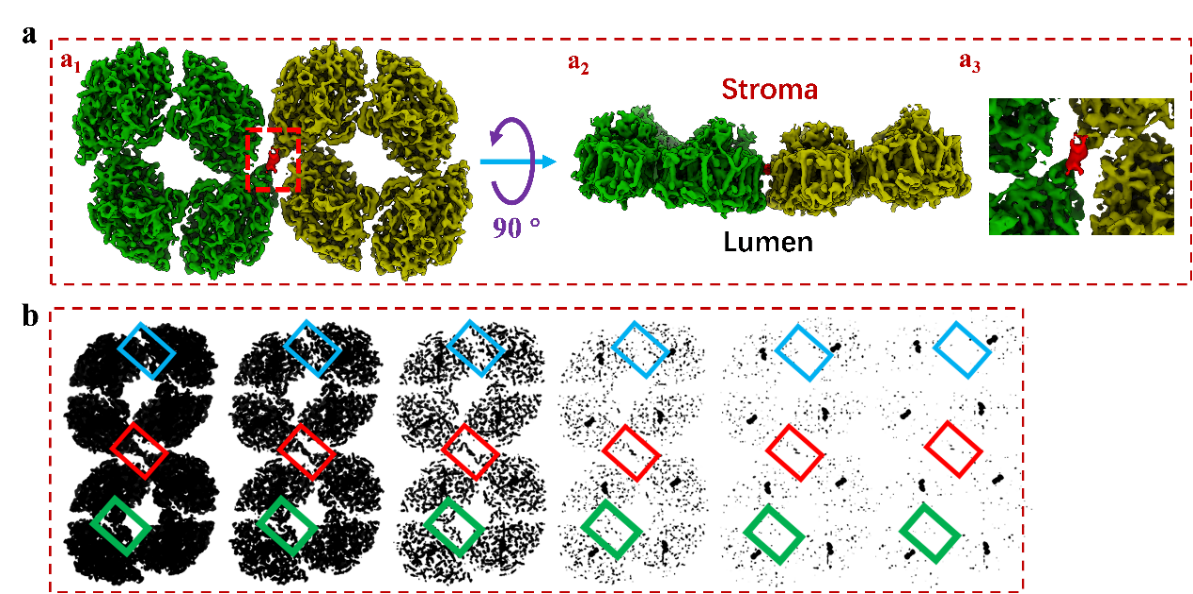

**Supplementary figure 4.** The linkage density in PSI octamer cryo-EM map. Top (a_1_) and side (a_2_) views of PSI octamer. The right up image shows the enlarge view of the red frame part, in which a interface for molecule connection is revealed. b-The electron density variation of the molecule interface in PSI dimer (green frame), tetramer (light blue frame) and octamer (red frame).

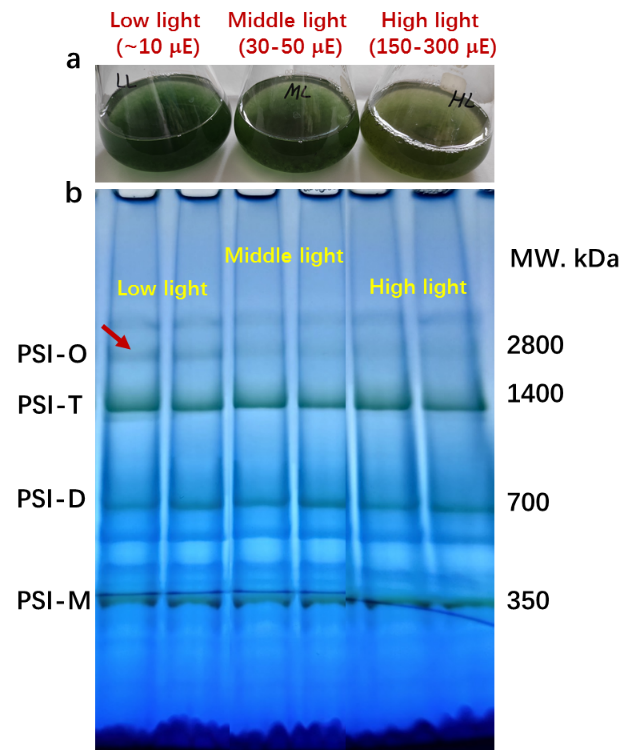

**Supplementary figure 5**. The PSI octamer response to the changing light conditions. Cell under log phase were incubated under low light, middle light and high light conditions for 48 hours. The content of PSI octamer was then detected biochemically with BN-PAGE experiment. There biological repeat experiments were proceeded to each group of samples.
